# Aerobic glycolysis supports hepatitis B virus biosynthesis through interaction between viral surface antigen and pyruvate kinase isoform M2

**DOI:** 10.1101/2020.08.10.244038

**Authors:** Yi-Hsuan Wu, Yi Yang, Ching-Hung Chen, Chia-Jen Hsiao, Tian-Neng Li, Kuan-Ju Liao, Bor-Sen Chen, Lily Hui-Ching Wang

## Abstract

As an intracellular pathogen, the reproduction of hepatitis B virus (HBV) depends on the occupancy of host metabolism machinery. Here we test a hypothesis if HBV may govern intracellular biosynthesis to achieve a productive reproduction. To test this hypothesis, we set up an affinity purification screen for host factors that interact with viral large surface antigen (LHBS). This identified pyruvate kinase isoform M2 (PKM2), a key regulator of glucose metabolism, as a binding partner of LHBS. We showed that the expression of LHBS affected oligomerization of PKM2 in hepatocytes, and thereby increased glucose consumption and lactate production, a phenomenon known as aerobic glycolysis. Interestingly, recovering PKM2 activity in hepatocytes by chemical activators, TEPP-46 or DASA-58, reduced biosynthesis of viral surface and core antigens. In addition, reduction of glycolysis by culturing in low-glucose condition or treatment with 2-deoxyglucose also decreased biosynthesis of viral surface antigen, without affecting general host proteins. Finally, TEPP-46 largely suppressed proliferation of LHBS-positive cells on 3-dimensional agarose plates, but showed no effect on the traditional 2-dimensional cell culture. Taken together, these results indicate that virus-induced metabolic switch may support *de novo* biosynthesis of HBV in hepatocytes. In addition, aerobic glycolysis is likely essential for LHBS-mediated oncogenesis. Accordingly, restriction of glucose metabolism may be considered as a novel strategy to restrain viral mediated biosynthesis and oncogenesis during chronic HBV infection.

**Author summary:** Chronic HBV infection is a life-long threat of patients, with a 25~40% increased risk of developing liver cirrhosis and cancer. Persistent expression of oncogenic viral products in the liver, especially LHBS, is an oncogenic caveat and resistant to current antiviral agents. Here we show that viral LHBS binds to host PKM2 and diminishes its kinase activity. This virus-host interaction induces metabolic switch from oxidative phosphorylation to aerobic glycolysis, with increased glucose consumption and lactate production. We show that such metabolic switch not only favors biosynthesis of HBV but also provokes hepatocarcinogenesis. Notably, restoration of PKM2 activity by chemical activators decreases expressions of viral products and largely suppresses virus-mediated hepatocarcinogenesis. This study highlights the importance of host metabolism in supporting viral biosynthesis and indicates a novel therapeutic approach to control chronic HBV infection via modulating host metabolic switch.

## Introduction

Hepatitis B is a life-threatening infectious liver disease caused by the hepatitis B virus (HBV). Chronic HBV infection affects ~292 million people worldwide (63.8% in Asia and 27.6% in Africa) in 2016, and has potential adverse outcomes that include hepatic decompensation, cirrhosis and/or hepatocellular carcinoma (HCC) [1]. WHO reported that 27 million people (10.5% of all people estimated to be living with chronic hepatitis B) were aware of their infection, while 4.5 million (16.7%) of the people diagnosed were on treatment (WHO. Global Hepatitis Report 2017. Geneva: 2017 ISBN: 978-92-4-156545-5). Though the currently available antiviral drugs can effectively reduce serum viral load in patients with chronic hepatitis B, complete elimination of the virus in the liver is still difficult. The blame goes to the long-lasting nature of an intracellular viral replication intermediate termed covalently closed circular (ccc) DNA, which serves as a viral persistence reservoir in the liver [2].

The HBV virion contains a compact 3.2-kb genome that exists as a partially double-stranded, relaxed circular DNA (rcDNA). Upon infection, cccDNA is generated as a plasmid-like episome in the host cell nucleus from the viral rcDNA genome [3]. Viral cccDNA is the template for all viral transcripts, and in consequence of new virions. The virion comprises an outer envelope of the lipid-embedded small (S), middle (M) and large (L) surface antigens (HBS) and an inner nucleocapsid (core particle; hepatitis B core antigen/HBcAg in serology). The three surface antigens collectively comprise HBsAg in serology. The virus also produces precore protein, serologically known as e antigen, or HBeAg. Both HBsAg and HBeAg play essential roles in chronic infection and are used as serological biomarkers of clinical examinations. High HBeAg is thought to induce T cell tolerance to HBeAg and HBcAg, which may contribute to early viral persistence upon initial infection [4]. Serum HBsAg have two resources, infectious virions enveloped with HBsAg and large amount of subviral particles consist of HBsAg. High HBsAg is believed to contribute to T cell exhaustion, resulting in limited or weak T cells response and even deletion of HBV-specific CD4 and CD8 T cells during T cell differentiation [5].

For patients with chronic hepatitis B, the treatment endpoint is the loss of HBsAg, known as “functional cure”, a state to indicate an effective control of HBV and long-term prognosis [6]. Spontaneous loss of HBsAg was detected in about 1.2% per year in treatment-naive patients in a recent systematic review and pooled meta-analyses reported [7]. It is noted that current antiviral therapies, such as type 1 interferons and nucleos(t)ide analogues (NUCs), rarely provoked the loss of HBsAg loss (0~10%), even after prolonged treatment [8]. As cccDNA is the intracellular viral reservoir, current pipelines of new HBV therapeutics have been focused on the eradication of the cccDNA [9]. On the other hand, several lines of evidence demonstrated that HBsAg was expressed not only from the cccDNA but also from the viral DNA integrated into the host genome. A recent RNAi-based treatment of chronically infected patients and chimpanzees also revealed that integrated hepatitis B virus DNA is a source of viral HBsAg, and even the dominant source in HBeAg-negative chimpanzees [10]. As HBV DNA integration may occur immediately upon infection [11], the abundance of hepatocytes with integrated HBV DNA in the liver may set a high threshold for the functional cure. Notably, integrated viral DNA with naturally derived PreS-truncation has been linked to the development of liver cancer. Several lines of evidence have showed that LHBS carrying PreS mutants, especially the PreS2 mutant LHBS, are driving factors of genomic instability and subsequent tumorigenesis [12–14]. A recent study reported that patients with PreS2 mutant were in high risk of hepatoma recurrence after curative hepatic resection [13]. Accordingly, persistence expression of intrahepatic LHBS is an oncogenic caveat in patients with chronic hepatitis B.

Because the loss of HBsAg is the indicator of endpoint treatment, there is interest in directly reducing expression of viral antigens and regulatory proteins. This study therefore aims to test if host metabolism may be tailored to control viral biosynthesis in the case of HBV. To this end, we set up to find intrahepatic metabolic regulators that may interact with HBsAg. Interestingly, protein pyruvate kinase isoform M2 (PKM2) is identified from the affinity purification with viral LHBS. PKM2 is a master regulator in glycolysis and has been implicated as a major metabolic switch in cancer metabolism [15]. Our evidence indicated that LHBS reduced PKM2 activity and thereby increased overall glucose consumption and lactate production in hepatocytes. Interestingly, modulation of glucose metabolism, either via chemical activation of PKM2 activity or the reduction of glucose supply, reduced HBV biosynthesis without affecting general host proteins. These data therefore indicate that intrahepatic viral biosynthesis may rely on the metabolic switch of host metabolism.

## Results

### PKM2 is a binding partner of LHBS

To explore the interaction between LHBS and host proteins, we set up an affinity purification experiment to identify potential intracellular binding partners of LHBS. Previously established stable cell lines carrying SNAP-tagged wild type LHBS [12, 16] were used for affinity purification and the resulted lysates were subjected to Mass-Spectrometry. PKM2 was identified in the SNAP-LHBS pulled-down lysates (data not shown). To confirm the interaction between PKM2 and LHBS, we performed a reciprocal immunoprecipitation. As shown in Fig 1A, PKM2 was detected in the pulled-down lysate of LHBS. In addition, LHBS was detected in the pulled-down lysate of PKM2. To understand which subcellular localization that LHBS and PKM2 interaction was taking place, we applied the proximity ligation assay (PLA) technology [17]. This technology provoked signal amplification of two complementary oligonucleotides that were linked to distinct antibodies specific for LHBS and PKM2. Using PLA technology, we showed that a subset of PKM2 interacted with LHBS mainly in the cytoplasm, but not in the cell nucleus (Fig 1B).

**Fig 1.**
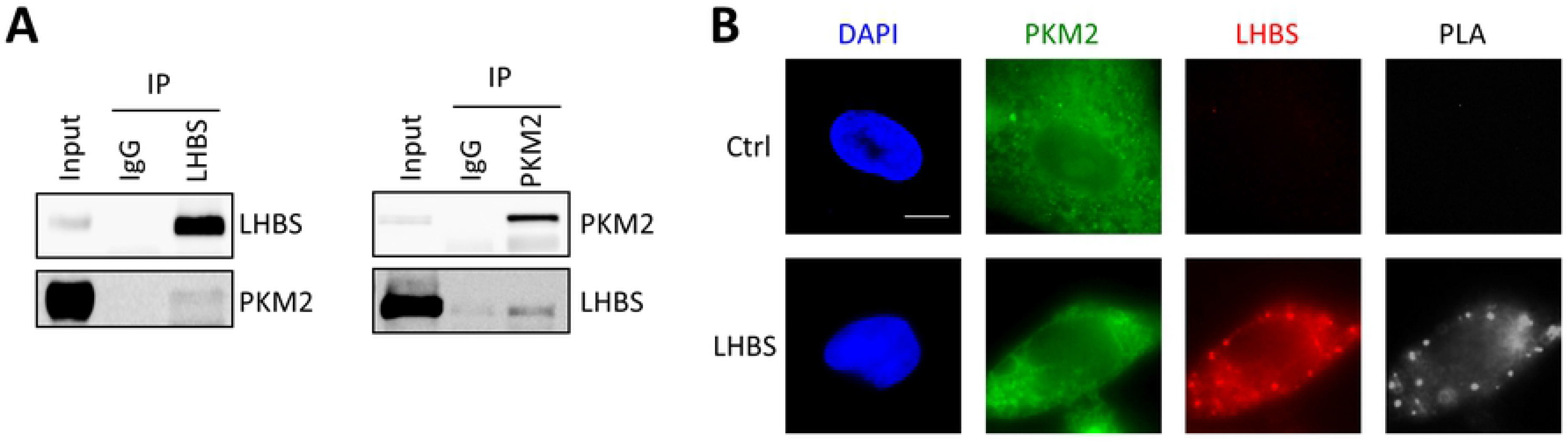
PKM2 is a binding partner of viral LHBS. (A) Reciprocal immunoprecipitation using control IgG and anti-LHBS or anti-PKM2 antibody was carried out using extracts prepared from a LHBS stable cell line. (B) The images showed subcellular localization between LHBS (red) and PKM2 (green) by proximity ligation assay (PLA, white). The bar indicates 5μm.

To identify essential interacting domains of PKM2 and LHBS, we made different expression constructs including SNAP-tagged LHBS (PreS1+S2+S), MHBS (PreS2+S), SHBS (S only), and PreS (PreS1+S2) (Fig 2A). Upon co-transfection with HA-tagged PKM2 into 293T cells, we detected LHBS, MHBS, and SHBS in the pull-down lysate of HA-PKM2, but not PreS, indicating that HBS was the major binding domain of PKM2 (Fig 2B). We next generated several constructs of PKM2 (Fig 2C) and performed transient transfection on a stable cell line expressing SNAP-LHBS (Fig 2D). Notably, whereas the full-length PKM2 weakly interacted with LHBS, HA-PKM2 110-531 showed a strong affinity with LHBS. This interaction was reduced upon serial truncation on the C-terminus, suggesting that C-terminus of PKM2 was important for binding to HBS. Finally, HA-PKM2 367-476 displayed a strong binding affinity with LHBS. Taken together, we mapped minimal interaction domains to the HBS region of LHBS and the C-terminus amino acid 367-476 of PKM2.

**Fig 2.**
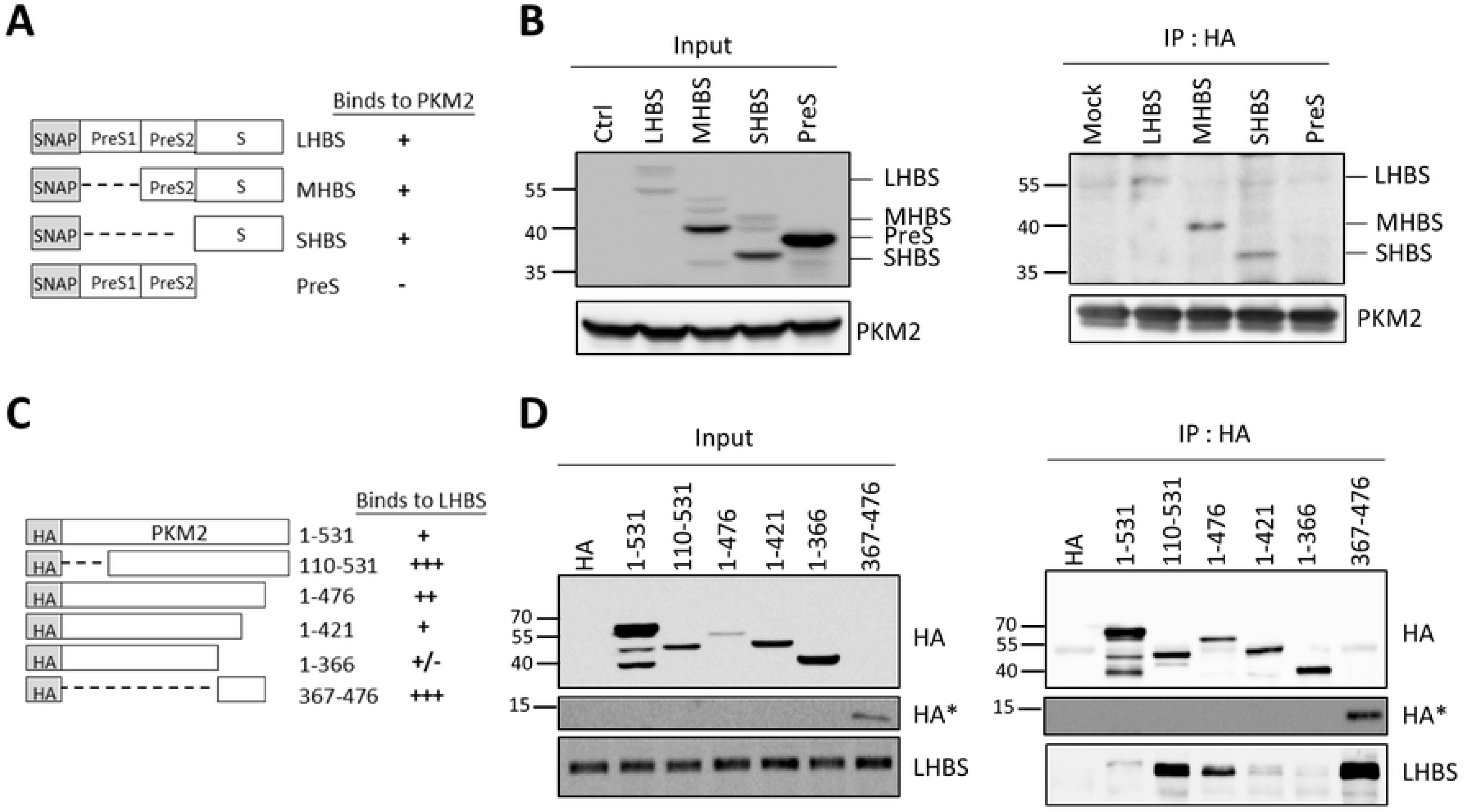
The minimal binding region between LHBS and PKM2. (A) All of the truncated LHBS fragments were ligated to SNAP tag. (B) Truncated LHBS fragments and HA-PKM2 were co-transfected in 293T cell line and pulled down HA, followed by blotting for SNAP-tag. (C) All of the truncated PKM2 fragments were ligated to HA tag. (D) Truncated PKM2 fragments were transfected in LHBS-expressing cells and pulled down HA, followed by blotting for HA-tag. *, High exposure image of low molecular weight proteins.

### LHBS provokes aerobic glycolysis via diminishing PKM2 activity

PKM2 catalyzes the last and physiologically irreversible step in glycolysis, the conversion of phosphoenolpyruvate (PEP) to pyruvate through the transfer of a phosphate group to ADP. We found that PKM2 activity was reduced by 86% in hepatocytes carrying LHBS (Fig 3A). As PKM2 exists as either a low-activity dimeric or high-activity tetrameric form [18], we next investigated whether LHBS may affect PKM2 oligomerization in a previously established LHBS stable cell line based on immortalized hepatocytes [16]. To this end, cell lysates were treated with 0.1% glutaraldehyde (GA) for crosslinking and then subjected to western blotting. In the control line, dimer form of PKM2 was detected after 10 minutes treatment with GA, and maximal at 20 minutes (S1 Fig). Similarly, dimer form of PKM2 was most abundant in LHBS-positive cells at 20 minutes post GA treatment. As over-crosslinking, defined by increased tetramerization, was detected in 25 and 30 minutes of GA treatment, we considered that 20 minutes of crosslinking was the best condition. Following 20 minute of GA treatment and in contrast to control cells, PKM2 dimerization was increased by 56% in LHBS cells (Fig 3B). Noted that overall PKM2 expression levels were equal in both control and LHBS cells, as shown by the same level of monomer in the absence of GA (Fig 3B, left). As dimeric PKM2 had lower kinase activity, the increased PKM2 dimer might explain the reduced kinase activity in LHBS cells.

**Fig 3.**
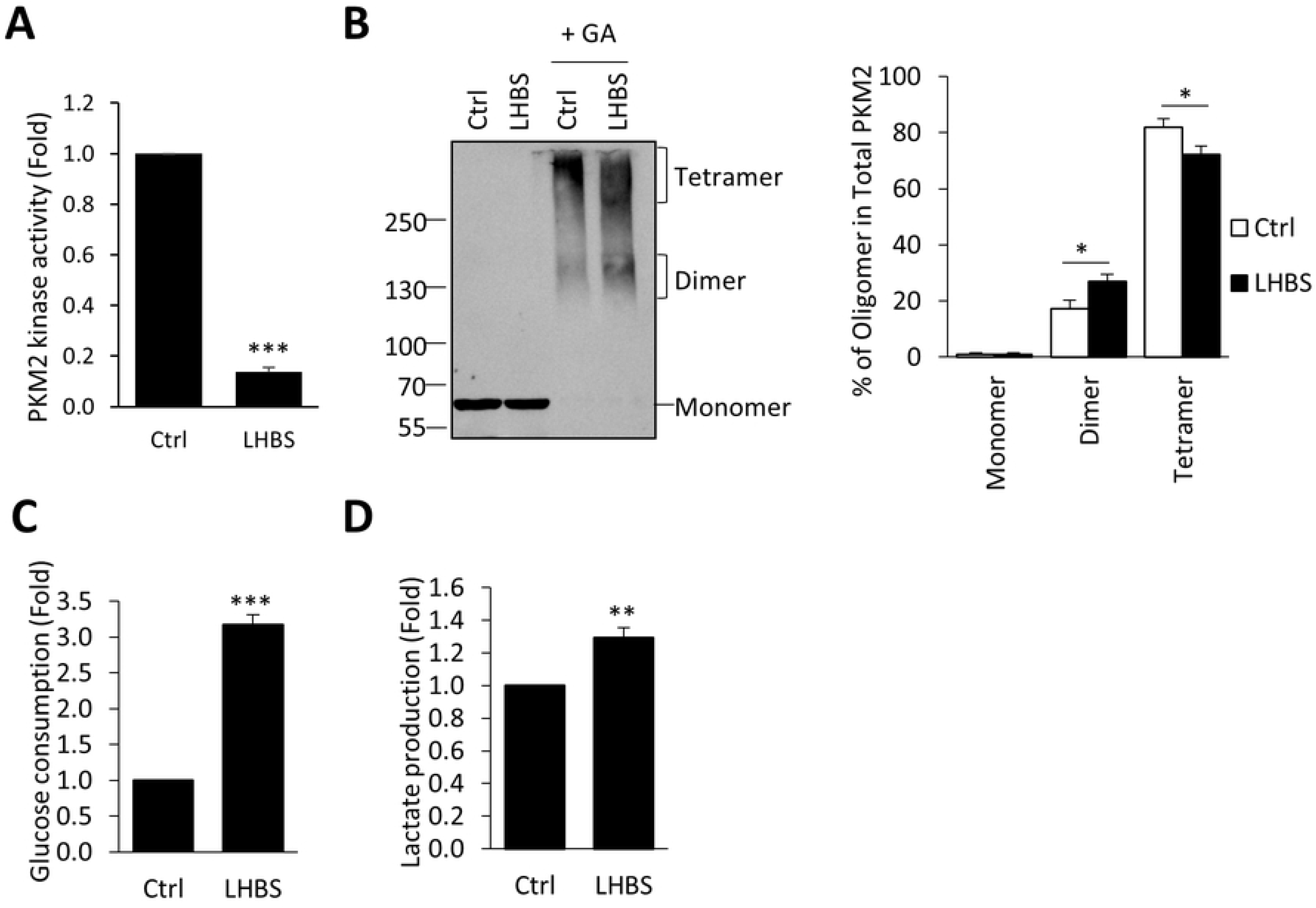
LHBS reduces PKM2 kinase activity and increases glucose consumption, as well as lactate production. (A) Pyruvate kinase activity of PKM2 was measured in control and LHBS-expressing cells. (B) PKM2 oligomerization was measured by cross-linking and western blotting. Control and LHBS-expressing cells were treated with GA for 20 minutes, followed by blotting for PKM2. The right panel showed quantitative result for the percentage of different PKM2 oligomers in control and LHBS cells. The culture media of control and LHBS cells were collected and subjected to measurements of glucose consumption (C) and lactate production (D). *, *p* <*0.05*; **, *P*<*0.01;* ***, *P*<*0.001*. Ctrl, control; GA, glutaraldehyde.

Reduction of PKM2 activity is necessary for the metabolic switch to aerobic glycolysis, also known as Warburg effect, which provides cancer cells with growth advantages [15]. Notably, glucose consumption was increased by 3.2 fold in LHBS cells (Fig 3C). In comparison to control cells, lactate production was increased by 29% in LHBS cells (Fig 3D). Taken together, LHBS reduced PKM2 activity and induced aerobic glycolysis in hepatocytes.

Although LHBS alone was sufficient to induce PKM2 dimerization, whether PKM2 activity is affected in the context of whole virus is not confirmed. Thus, we used recombinant adenovirus carrying wild type HBV (Ad-HBV-WT) or control knockout (Ad-HBV-KO) genome as an *in vitro* model [19] and examined the effect of PKM2 upon infection. We infected HuH-7 cells with Ad-HBV-WT or Ad-HBV-KO or a 1:1 mixture in 200 MOI for 2 days and examined the expression of LHBS and PKM2 oligomerization. As expected, LHBS expression was positively correlated with the dose of Ad-HBV-WT (Fig 4A). In addition, PKM2 dimerization increased upon infection with Ad-HBV-WT, in a dose-dependent manner (Fig 4B and 4C). Similar results were observed in HepG2 cells infected with Ad-HBV-WT, except that 100 MOI was used for HepG2 (Fig 4D-4F). Finally, we confirmed that kinase activity of PKM2 was reduced by 13% in HuH-7 cells infected with Ad-HBV-WT (S2 Fig). These data confirmed that aerobic glycolysis could be induced by HBV.

**Fig 4.**
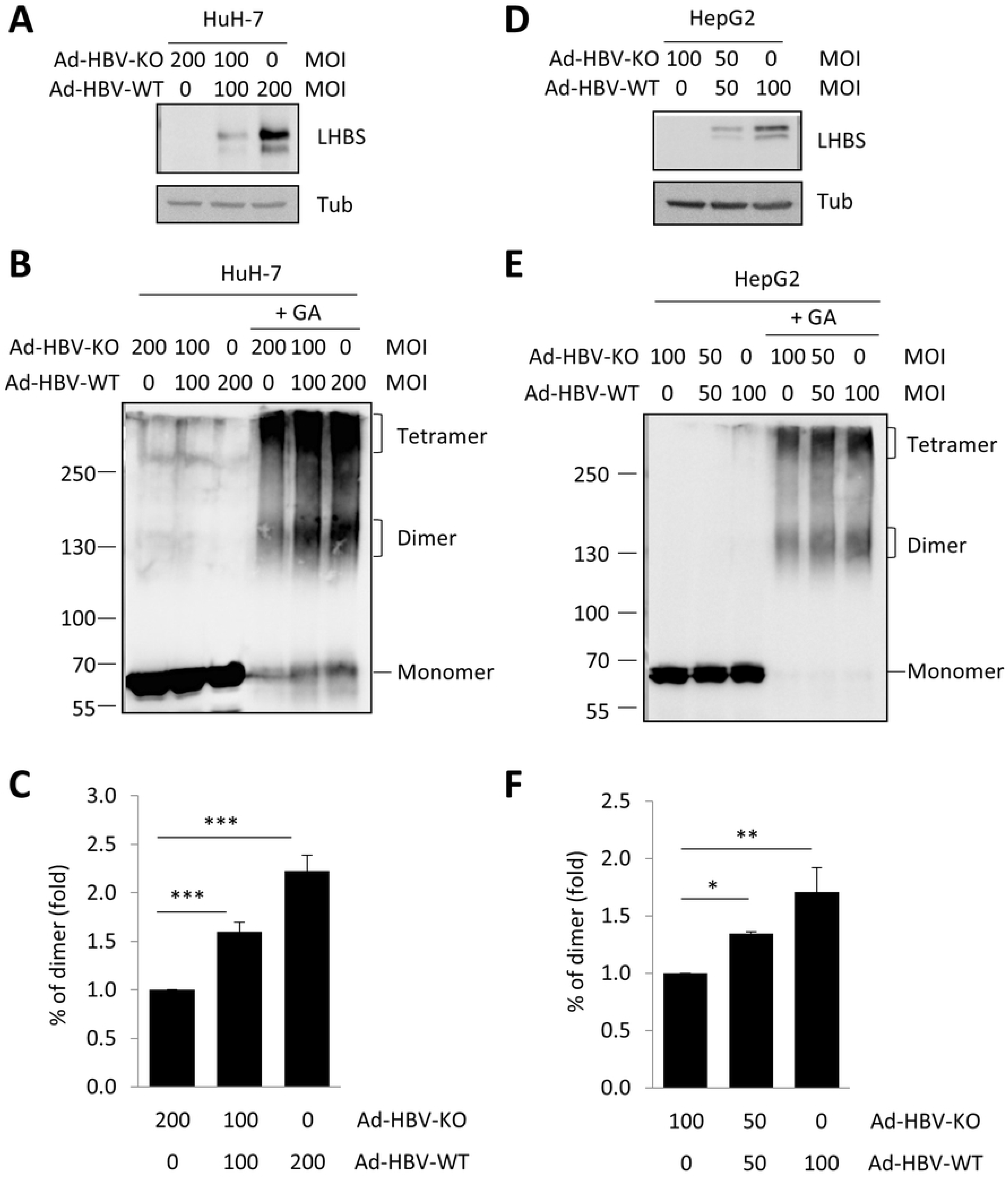
Infection of Ad-HBV-WT increases PKM2 dimerization. (A) HuH-7 cells were infected with either Ad-HBV-KO or Ad-HBV-WT for 2 days at different multiplicity of infection (MOI). Expressions of viral LHBS and Tubulin were shown. (B) The Ad-HBV infected HuH-7 cells were treated with either PBS or GA for 20 min and then subjected to Western blotting analyzing PKM2 oligomerization. Changes in the percentage of PKM2 dimer was shown in (C). (D) HepG2 cells were infected with either Ad-HBV-KO or Ad-HBV-WT for 2 days at different MOI. Expressions of viral LHBS and Tubulin were shown. (E) The Ad-HBV infected HepG2 cells were treated with either PBS or GA for 20 min and then subjected to Western blotting analyzing PKM2 oligomerization. Changes in the percentage of PKM2 dimer was shown in (F). *, *P*<*0.05;* **, *P*<*0.01;* ***, *P*<*0.001*. KO, knockout; WT, wildtype; GA, glutaraldehyde.

### PKM2 negatively regulates biosynthesis of HBV

As an intracellular pathogen, viruses may hijack host metabolism to support their own propagation. Specifically, viral infection may trigger metabolic reprogramming in host cells to facilitate optimal virus production [20]. We therefore suggested that the metabolic switch induced by LHBS might be essential for HBV biosynthesis. To test this hypothesis, we asked if activation of PKM2 activity may affect biosynthesis of HBV in the host. Specifically, we transiently transfected pHBV3.6 plasmid into HuH-7 cells and treated with TEPP-46, a compound known to stabilize tetrameric PKM2 [21]. Upon treatment with TEPP-46, extracellular secretions of HBsAg and HBeAg were reduced by 40% and 20%, respectively (Fig 5A). In addition, intracellular expressions of LHBS, SHBS, and HBcAg were reduced by 30, 41, and 26%, respectively (Fig 5B and 5C). In contrast, expression levels of viral HBx and general host proteins, including GRP78, HSC70, PKM2, Plk1, Mad2L1, Bcl2, and PCNA, were not affected by TEPP-46.

**Fig 5.**
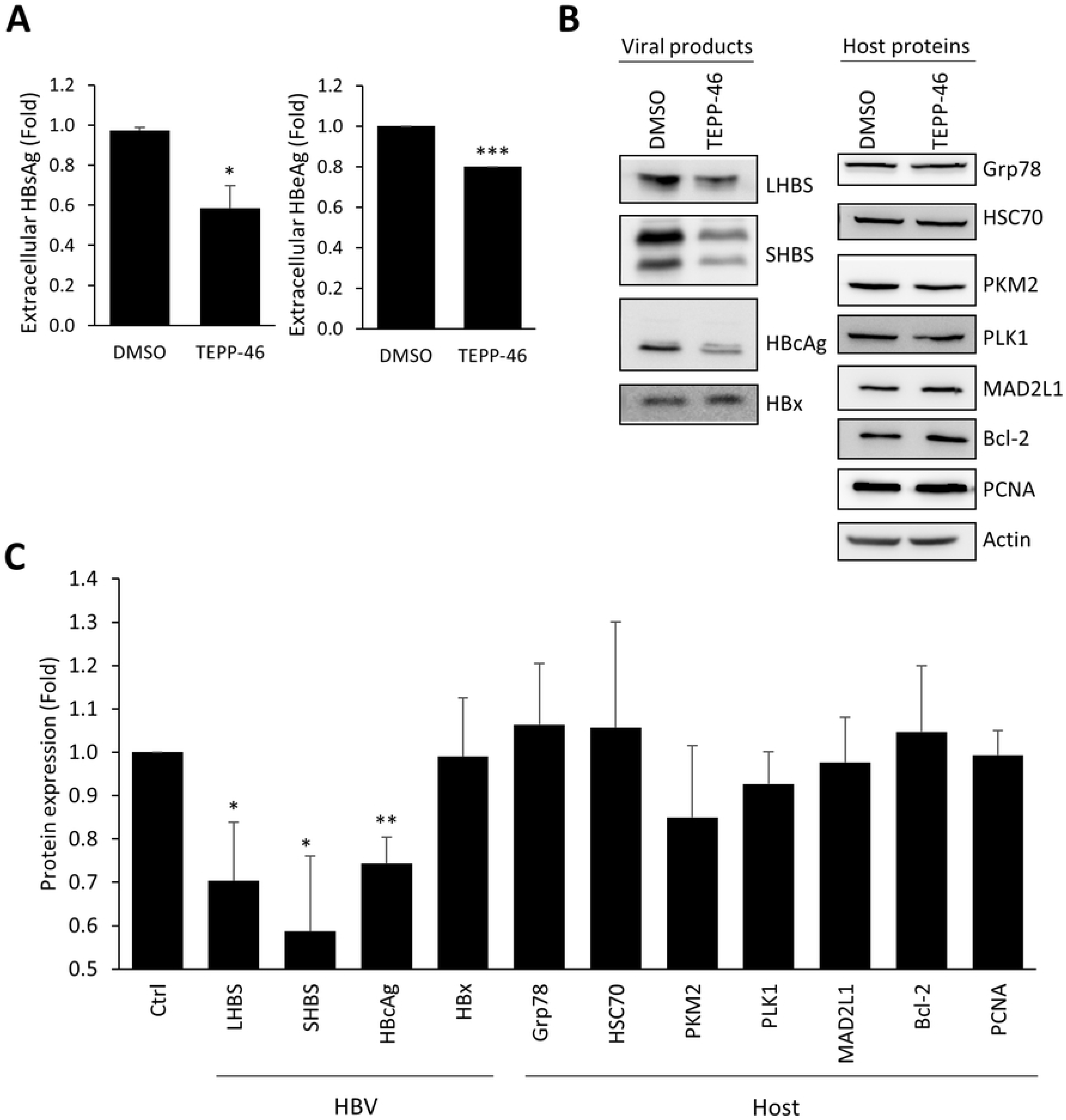
TEPP-46 reduces biogenesis of HBV in HuH-7 cells. HuH-7 cells were transfected pHBV3.6 for 24 hr and then treated with DMSO and 20 μM TEPP-46 for additional 24 hr. (A) The culture media of both treatment were collected and subjected for measurement of HBsAg and HBeAg level. (B) Expressions of HBV viral products (left panel) and host proteins (right panel) upon treatment of DMSO or TEPP-46 were shown. (C) Statistic result of viral and host protein expression was shown. *, *P*<*0.05;* **, *P*<*0.01;* ***, *P*<*0.001*.

Similar results were obtained in cells treated with DASA-58, another PKM2 activator [21]. PKM2 activity in LHBS cells was increased by DASA-58 (Fig 6A). As expected, extracellular secretions of HBsAg and HBeAg were also reduced (Fig 6B). We confirmed that intracellular expressions of viral LHBS, SHBS, and HBcAg were reduced upon DASA-58 treatment, except for viral HBx (Fig 6C and 6D). Finally, we asked if restoration of PKM2 activity may reduce intermediate metabolites of pentose phosphate pathway. This was confirmed by the reduction of 6-phosphogluconic acid (6-PGA) in cells treated with TEPP-46 and DASA-58 (Fig 6E). Taken together, these data indicated that restoration of PKM2 activity might suppress intracellular biosynthesis HBV, likely due to the lack of supporting intermediate metabolites from glycolysis.

**Fig 6.**
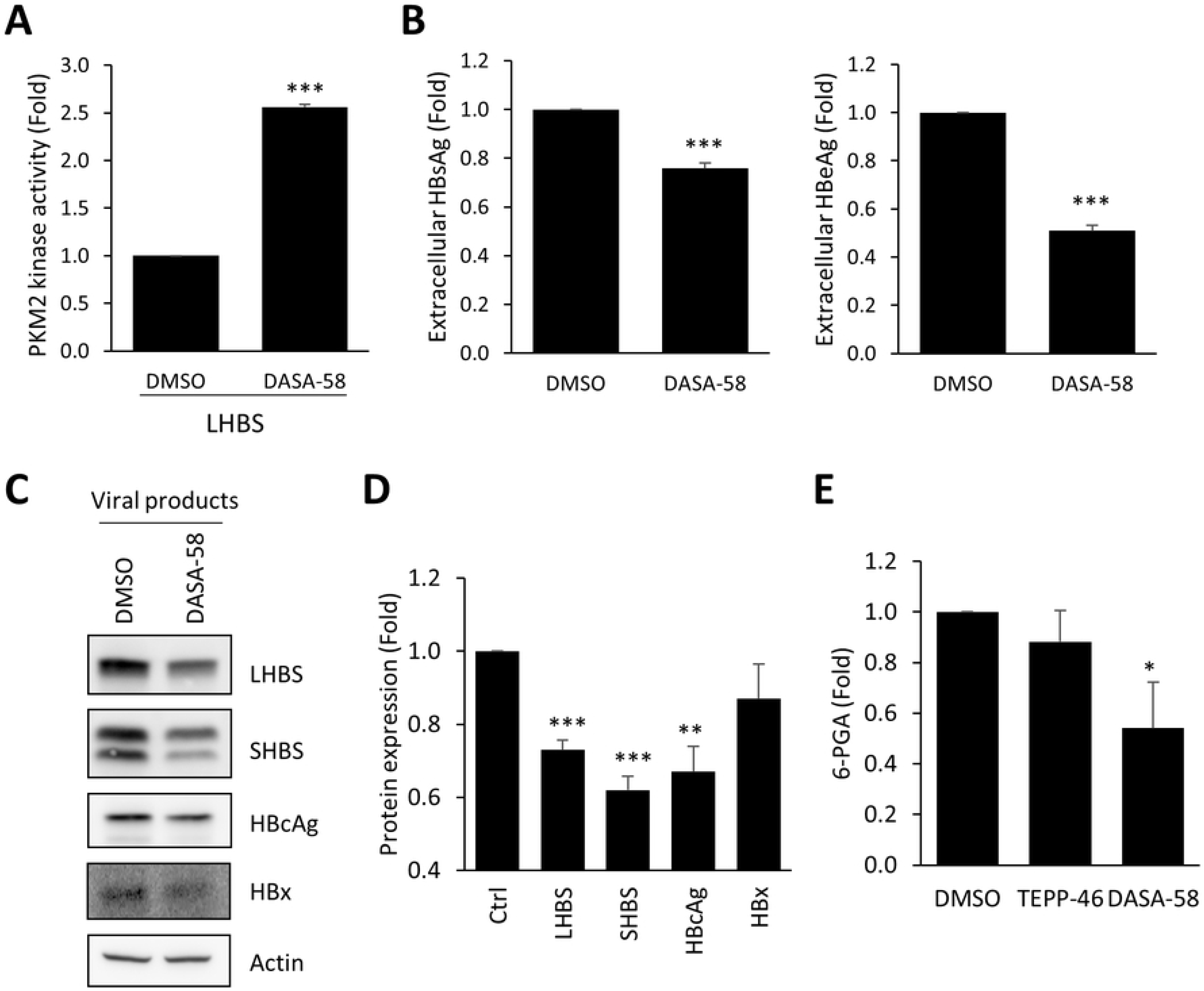
DASA-58 reduces viral biogenesis. (A) DMSO and 50 μM DASA-58 were treated in LHBS-positive cells and lysates were collected to perform pyruvate kinase activity assay. (B) HuH-7 cells were transfected pHBV3.6 for 24hr and then treated with DMSO and 50 μM DASA-58 for additional 24 hr. The culture media were collected for measuring HBsAg and HBeAg level. (C) Expressions of HBV viral products upon treatment of DMSO or DASA-58 were shown. (D) Statistic result of viral protein expression was shown (E) The concentration of 6-PGA was measured in HuH-7 cells. *, *P*<*0.05;* **, *P*<*0.01;* ***, *P*<*0.001*.

### Productive biosynthesis of HBV relies on aerobic glycolysis

Next we asked if HBV biosynthesis can be regulated by modulating glucose metabolism. We found that extracellular secretions of HBsAg and HBeAg were both reduced in low glucose condition without affecting general cell viability (Fig 7A and 7B). Interestingly, we only observed reduction of intracellular LHBS under low glucose condition but not HBcAg (Fig 7C). Accordingly, we suspected that biosynthesis of HBsAg was more sensitive to the glucose supply than that of HBcAg.

**Fig 7.**
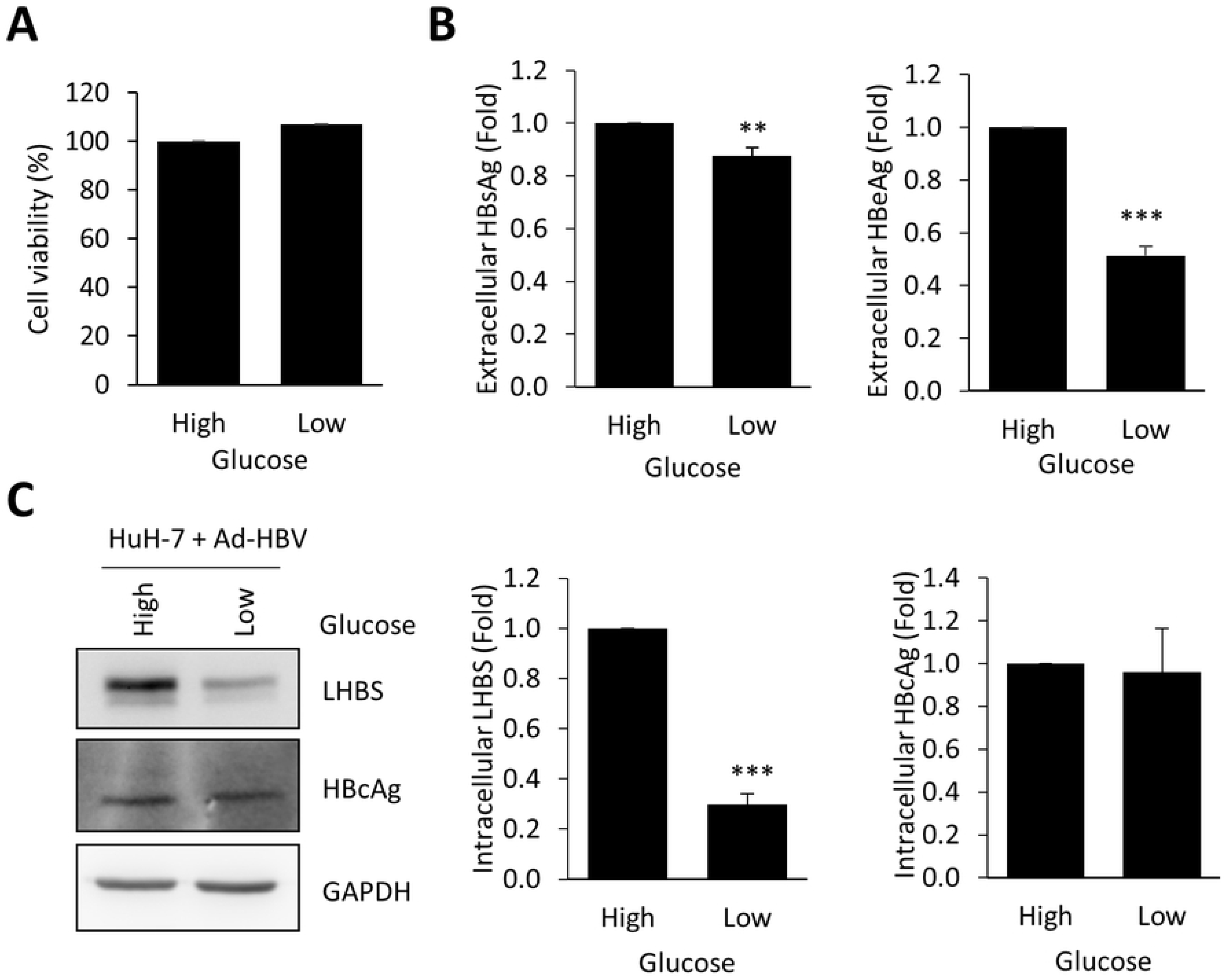
Transient low glucose treatment reduces biogenesis of HBV. (A and B) HuH-7 cells were infected with Ad-HBV-WT for 24 hr and then treated with medium containing high (4.5g/L) or low (1g/L) glucose for additional 24 hr. (A) The cells were collected for measuring cell viability. (B) The culture media were collected for measuring HBsAg and HBeAg level. (C) Left panel showed protein expression of viral proteins in HuH-7 infected cells. Middle and right panels showed statistic result of viral protein expression. **, *P*<*0.01;* ***, *P*<*0.001*.

2-Deoxy-D-Glucose (2-DG) is a glucose molecule which has the 2-hydroxyl group replaced by hydrogen, so that it acts to competitively inhibit the production of glucose-6-phosphate from glucose by phosphoglucoisomerase, thereby interrupting glycolysis. Here we showed that both glucose consumption and lactate production were suppressed by 2-DG in a dose-dependent manner (Fig 8A). In addition, extracellular secretions of HBsAg and HBeAg were reduced by 2-DG in a dose-dependent manner (Fig 8B). Coordinately, intracellular LHBS and HBcAg were reduced by 2-DG (Fig 8C and 8D). Taken together, our data indicated that major HBV biosynthesis, especially HBsAg and HBcAg, could be inhibited by blocking glycolysis using 2-DG.

**Fig 8.**
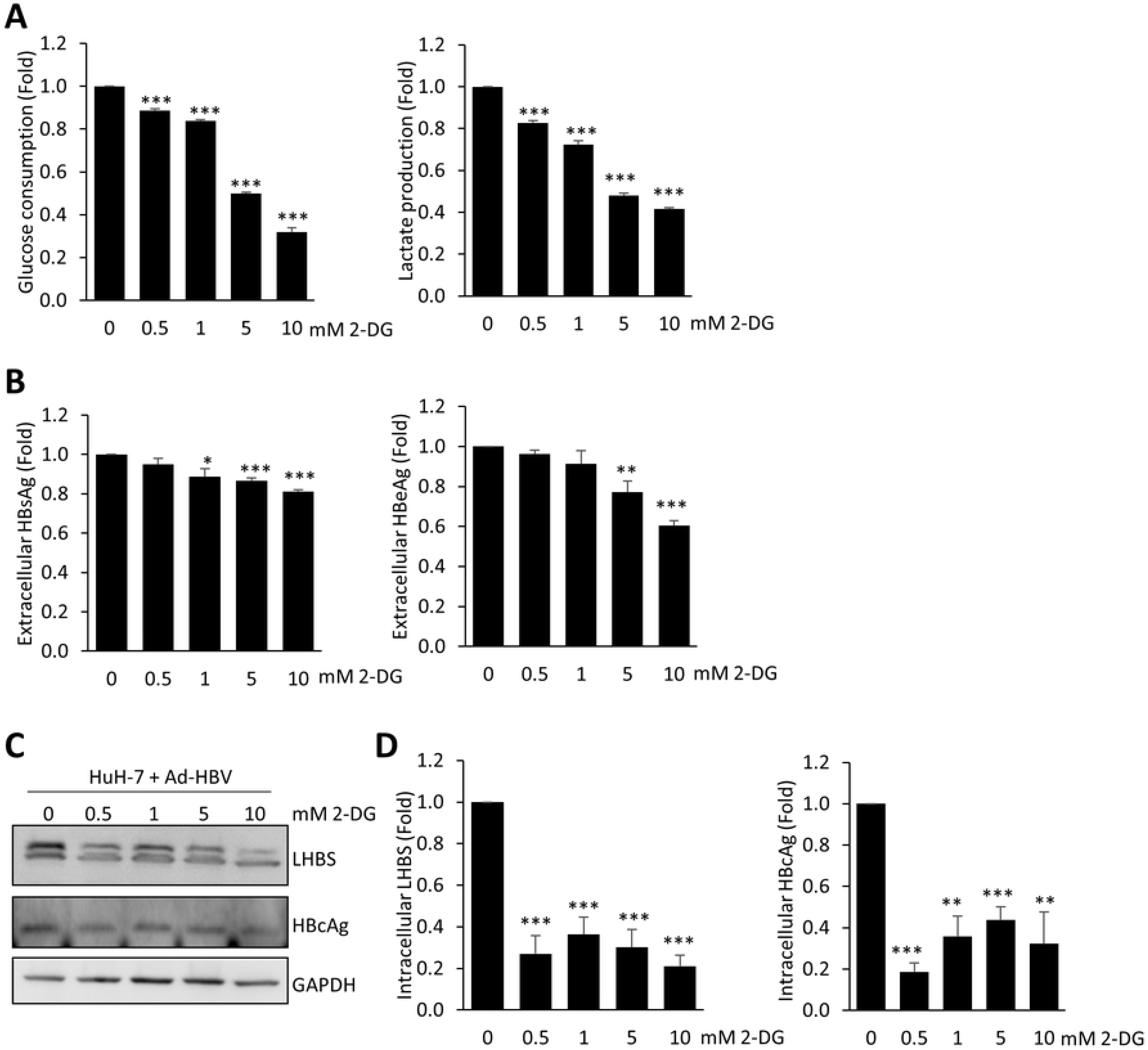
Low dose of 2-DG reduces aerobic glycolysis and viral protein expression. HuH-7 cells were infected with Ad-HBV-WT for 24 hr and then treated with different dosages of 2-DG for another 24 hr. (A) The media were collected for measuring glucose consumption (left) and lactate production (right). (B) Extracellular HBsAg and HBeAg level were analyzed from the media. (C) Protein expression of LHBS and HBcAg were verified 48 hours after infection and 2-DG treatment. (D) Statistic result of protein expression of LHBS and HBcAg was shown. *, *p* <*0.05;* **, *P*<*0.01;* ***, *P*<*0.001*.

### Activators of PKM2 suppress LHBS-mediated oncogenesis

Intrahepatic LHBS has been reported as a priming factor for the development of HCC [22]. The expression of LHBS induced cytokinesis failure and consequent aneuploidy via induction of DNA damage and polo-like kinase 1 (PLK1)-mediated G2/M checkpoint failure in hepatocytes [12]. Whether aerobic glycolysis involved in LHBS-mediated oncogenesis is yet to be determined. Here we showed that LHBS induced anchorage-independent growth of immortalized hepatocytes on the 3-dimensional condition of soft agar plates. In contrast, the control lines failed to grow on soft agar plates (Fig 9A). We showed that TEPP-46 restored PKM2 activity (Fig 9B) without affecting general cell viability under traditional 2-dimensional culture (Fig 9C). However, cell growth on soft agar plates was largely diminished by TEPP-46 (Fig 9D and 9E), as well as DASA-58 (Fig 9F and 9G). Taken together, treatments of PKM2 activators could largely suppressed LHBS-mediated anchorage-independent growth in hepatocytes. These data therefore provided a strong evidence to support the role of PKM2 as a negative regulator of HBV-mediate hepatocarcinogenesis.

**Fig 9.**
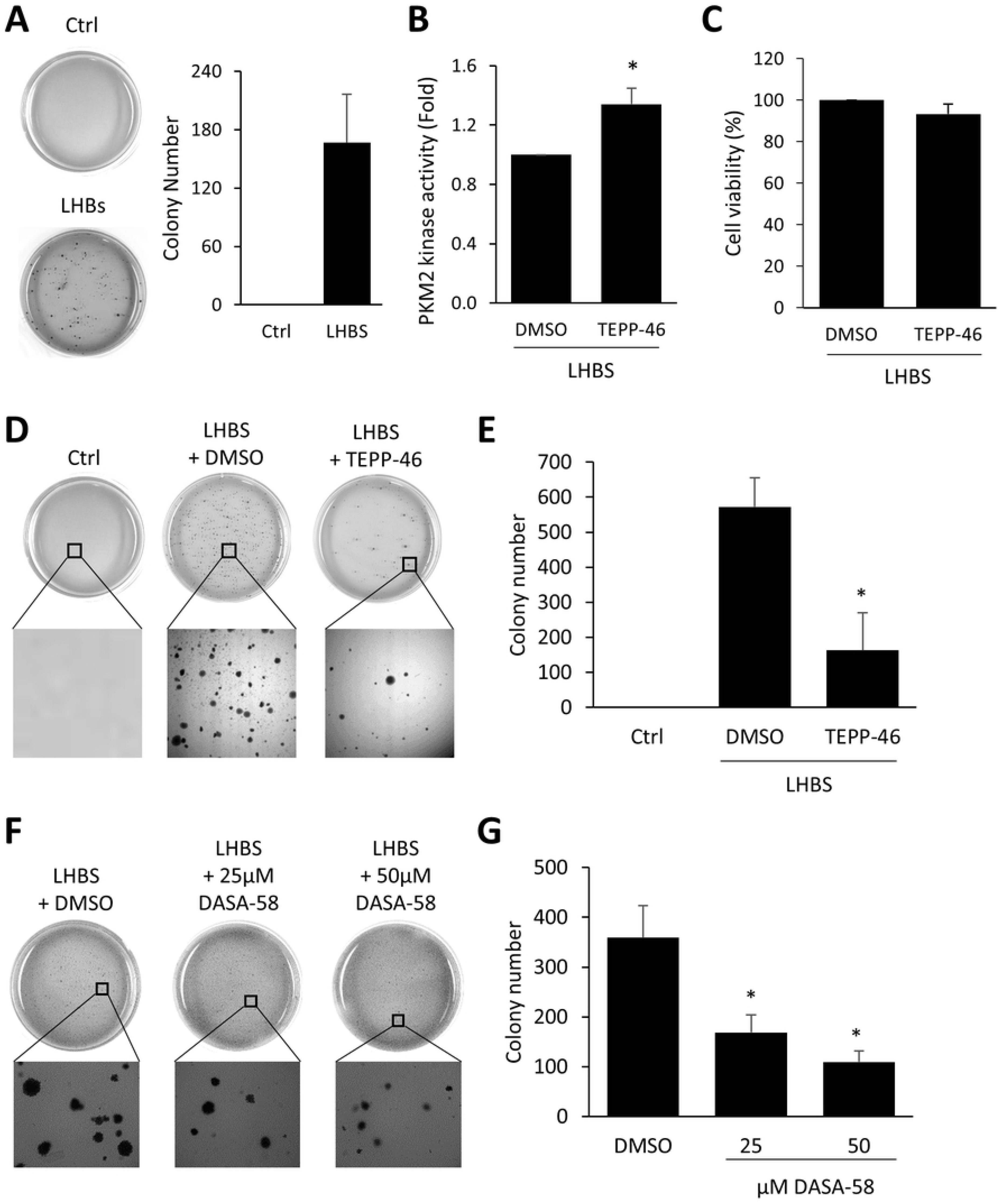
PKM2 activators suppressed LHBS-mediated oncogenesis. (A) Colony number was checked between control and LHBS-positive cells. (B) LHBS-positive cells were collected 24 hours after 20 μM TEPP-46 treatment to measure PKM2 kinase activity. (C) Cell viability was measured 48 hours after TEPP-46 treatment in LHBS-positive cells. (D) Representative colony result was shown. (E) Statistic result of colony number between control and TEPP-46 treated LHBS-positive cells was shown. (F) Colony number was checked between treatment of DMSO and DASA-58 in LHBS-positive cells. Representative colony result was shown. (G) Statistic result of colony number between control and DASA-58 treated LHBS-positive cells was shown. *, *p* <*0.05*. Ctrl, control.

## Discussion

Viral hijacking of cellular metabolism has been noted for over half a century. Pushing the metabolic flux into biosynthetic pathways is the key to support *de novo* synthesis of viral building blocks. The mechanisms and consequences of virus-induced metabolic switch, or reprogramming, have only begun to be studied in detail over the past decade [20]. This study adds a new model mechanism to illustrate how glucose metabolism is switched to aerobic glycolysis through the interaction between a key metabolic enzyme and a viral product, in this case, PKM2 and HBV LHBS. Noted that metabolic phenotypes conferred by viruses often imitate metabolic changes in cancer cells. As such, increased glucose consumption and lactate production are two major phenotypes of Warburg effect, a well-known cancer metabolism that depicts increased glycolysis in the presence of oxygen. In cancer cells, such metabolic reprogramming is likely beneficial, as it allows rapid biosynthesis to support growth and proliferation. Aerobic glycolysis also enhances disruption of tissue architecture and immune cell evasion in the tumor microenvironment [23]. As for virus, the most obvious beneficial effect comes from the flux into biosynthetic pathways, which support the *de novo* synthesis of viral building blocks [20]. In this study, we find that HBV biosynthesis, especially HBsAg, relies on the metabolic switch regulated by PKM2. Interestingly, chemical activators of PKM2 only suppressed major biosynthesis of HBV but not general host machinery (Fig 5). Our results indicated that viral biosynthesis is more sensitive to metabolic rewiring than host biogenesis. As such, these results imply that chronic HBV infection can be controlled by rewiring the metabolic switch. If so, targeting host metabolic switch can be a promising approach to control viral replication during chronic HBV infection.

Current study did not address why HBV biosynthesis is more sensitive to metabolic rewiring than the host machinery. In this study, we show that expressions of 8 different host proteins, including mitotic and non-mitotic proteins, were not affected by TEPP-46, whereas, most viral products were reduced in response to TEPP-46 (Fig 5B). We suspect that HBV biosynthesis may rely on *de novo* synthesis with newly synthesized building blocks, probably come from products of pentose phosphate pathway. This notion is partly supported by the observation that 6-PGA, a product of pentose phosphate pathway, was reduced upon PKM2 activation (Fig 6E). In addition, we suspect that the impact of PKM2 activator on HBV biosynthesis is regulated at the translational level, but not the transcription, as HBV RNA was not affected upon PKM2 activation (data not shown).

In addition to support viral biosynthesis, increased glucose consumption in HBV-infected cells also imply that these cells have a great need for energy, similar to highly proliferating cells or cancer cells. Interestingly, general viability of LHBS cells were not affected by TEPP-46 or DASA-58 when cultured in traditional two-dimensional cell culture, but were largely suppressed when cultured in soft-agar plates (Fig 9). These results indicate that aerobic glycolysis is likely irrelevant under two-dimensional culture, but is essential for cell proliferation under three-dimensional culture condition. Accordingly, treatments with PKM2 activators may have two beneficial effects on patients with chronic HBV infection: to suppress viral biosynthesis and to prevent proliferation of virus-infected hepatocytes. It has been shown that TEPP-46 has a good oral bioavailability *in vivo* and showed a promising effect in reducing tumor size in lung cancer xenograft tumors [21]. Accordingly, we suggest that TEPP-46 may be considered as a chemoprevention agent for patients with chronic HBV infection.

In this study, we also found that low glucose condition limited HBV biosynthesis in hepatocytes, implying that glucose support may affect viral reproduction. Notably, chronic hepatitis B patients with diabetes were shown to have 2.3-fold increased risk of developing HCC [24]. The link between serum glucose level and HBV replication is not clarified at this stage. It will be interesting to learn if controlling blood sugar may prevent HCC development in chronic hepatitis B patients with sustained diabetes. In this case, anti-diabetes agents may be applied to control HBV biosynthesis in patients with diabetes. On the other hand, we don’t know whether anti-diabetic agents, such as metformin and alpha-glucosidase inhibitors, can be used in patients without diabetes and whether anti-diabetic agents not leading to hypoglycemia are still beneficial in such condition. These questions are awaiting for further investigations.

In summary, our data provide a mechanistic view to explain how metabolic switch is induced upon chronic HBV infection and thereby to support biosynthesis of HBV. This study suggest that rewiring the host metabolic switch may be applied to control viral biosynthesis and hepatocarcinogenesis in patients with chronic hepatitis B.

## Materials and methods

### Cell culture and transfection

The hTERT-immortalized hepatic progenitor cell line NeHepLxHT was obtained from the American Type Culture Collection (ATCC, Manassas, VA, USA). Stable cell lines expressing SNAP tag (as a control) or SNAP - tagged LHBS were established from NeHepLxHT cells that have been described previously [16]. Both of the stable cell lines were cultured on type-I collagen dish in Dulbecco’s modified of Eagle’s medium/Ham’s F-12 50/50 mix (DMEM-F12) supplemented with 15% fetal bovine serum (FBS) (Biological industries), 1x penicillin-streptomycin (Corning), ITS premix (5 μg/ml insulin, 5 μg/ml human transferrin, and 5 ng/ml selenic acid; BD), 20 ng/ml epidermal growth factor (BD), and 100 nM dexamethasone (sigma). 293T, HepG2 and HuH-7 cells were cultured in standard DMEM-based media. All the cell lines incubated at 37 °C in a humidified atmosphere containing 5% CO_2_. DNA transfection was performed with GeneJet In Vitro DNA Transfection Reagent (Ver. II) (SignaGen Laboratories) according to the manufacturer’s instructions.

### Infection with Ad-HBV

Productions of Ad-HBV-KO and Ad-HBV-WT were followed the previous study [19], kindly provided by Dr. Li-Rung Huang (National Health Research Institutes, Taiwan). Adenoviral vector stocks was 10^11^ infectious units (i.u.)/ml. Briefly, HuH-7 and HepG2 cells were seeded in 6 cm dishes for overnight. Infected the cells with Ad-HBV-KO or Ad-HBV-WT at a multiplicity of infection (MOI) of 0 to 200 genome equivalents/cell for 2 days. For indicated drug treatments, cells were infected with Ad-HBV for 24 hour, followed by treatments with indicated drugs for another 24 hour.

### Reagents

TEPP-46 (ML-265) and 2-deoxy-D-Glucose (2-DG) were ordered from Cayman Chemical. DASA-58 was purchased from SelleckChem. Dimethyl sulfoxide (DMSO) was used as vehicle control. All inhibitor stocks were made in DMSO with the following stocking concentrations: TEPP-46, 20 mM; 2-DG, 1 M; DASA-58, 50 mM.

### Antibodies

The following antibodies were used in this study: rabbit anti-PKM2 (CST 4053, Cell Signaling Technology), rabbit anti-HBcAg (B0586, DAKO), mouse anti-HA-tag (sc-7392, Santa Cruz), mouse anti-HSC70 (sc-7298, Santa Cruz), rabbit anti-SNAP-tag (P9310, New England BioLabs), rabbit anti-beta-tubulin (NB600-936, Novus Biologicals), mouse anti-beta-actin (NB600-501, Novus Biologicals), rabbit anti-GAPDH (GTX100118, GeneTex), rabbit anti-Grp78 (GTX113340, GeneTex), rabbit anti-PLK1 (GTX104302, GeneTex), rabbit anti-MAD2L1 (GTX104680, GeneTex), rabbit anti-Bcl-2 (GTX100064, GeneTex), and rabbit anti-PCNA (GTX100539, GeneTex), peroxidase-conjugated AffiniPure goat anti-rabbit IgG (111-035-003, Jackson ImmunoResearch), peroxidase-conjugated AffiniPure goat anti-mouse IgG (115-035-003, Jackson ImmunoResearch). Mouse anti-LHBS (7H11), mouse anti-SHBS (86H6), and mouse anti-HBx (20F3) were kindly provided by Ning-Shao Xia (Xiamen University, China).

### Western blot analysis

Cells were lysed in RIPA buffer (50 mM Tris pH7.5, 150 mM Sodium chloride, 1% Nonidet P40 Substitute, 0.5% Sodium deoxycholate, and 0.1% SDS) containing one protease inhibitor cocktail (Roche). Centrifuged at full speed at 4°C for 15 minutes then collected supernatant and measured protein concentration by Bradford assay. Added 2x laemmli buffer (0.04% SDS, 20% glycerol, 120mM Tris pH6.8) containing β-ME and boiled at 100 °C for 15 minutes. 30 μg of total protein was analyzed by SDS-PAGE gel electrophoresis and transferred to a PVDF membrane (Millipore). The membrane was probed with antibodies at 4 °C overnight and visualized using Western Lightning™ Plus ECL (erkinElmer. Inc).

### Immunoprecipitation

Different SNAP-tagged truncated LHBS constructs (SNAP as a control, SNAP-LHBS, SNAP-MHBS, SNAP-SHBS, and SNAP-PreS) and HA-tagged PKM2 were transfected into 293T cells or different HA-tagged truncated PKM2 constructs (full length, 110-531 a.a., 1-476 a.a., 1-421 a.a., 1-366 a.a., and 367-476 a.a.) were transfected into LHBS stable cells. Cells were treated with RIPA buffer (50 mM Tris pH7.5, 150 mM Sodium chloride, 1 mM EGTA, 0.5% Nonidet P40 Substitute, 0.1% Sodium deoxycholate) containing one protease inhibitor cocktail and 1mM DTT. Took 40μl protein G Mag Sepharose (GE Healthcare) and incubated with antibody at 4°C for 30 minutes. Collected the cell lysate and measured protein concentration, diluted protein to 2-5 μg/μl. Took part of cell lysate into new tube and added equal volume of 2x laemmli buffer, boiled at 100°C for 15 minutes (Input). The rest of cell lysate were added into antibody-Mag sepharose mixture. Rocked at 4°C for overnight then added 2x laemmli buffer, boiled at 100°C for 15 minutes (IP product). Samples were detected by Western blotting.

### Proximity ligation assay

Cells were seeded on collagen-coated coverslips and washed with PBS then fixed with 4% formaldehyde/PBS at room temperature for 10 minutes. Incubated the primary antibodies for one hour then applied two PLA probes at 37°C for one hour. Next, added ligase and ligation buffer for 30 minutes then added polymerase and amplification buffer at 37°C for 3 hours. Finally, incubated with AlexaFluor-conjugated secondary antibody for 1 hour and mounted the coverslip with DAPI-containing mounting medium. The fluorescence signal was detected by Leica DMI6000 inverted microscope.

### PKM2 kinase activity assay

The assay followed the protocol of Pyruvate Kinase Activity Colorimetric/Fluorometric Assay Kit (BioVision). The cells was extracted with 4 volumes of assay buffer then centrifuged to get clear extract. Added 50 μl diluted sample into 96 well plate then 50 μl reaction mix for each well. Measured OD 570 nm at T1 to read A1. Measured again at T2 after incubating at 25°C for 10-20 minutes to read A2. PK activity was calculated against the standard curve.

### Measuring glucose consumption and lactate production

Cells were cultured on 96-well plate for 3 days and the supernatant was collected. Glucose consumption was analyzed using glucose assay kit (Eton Bioscience), according to manufacturer’s protocol. Lactate production was analyzed using lactate assay kit (Eton Bioscience), followed the manufacturer’s protocol.

### PKM2 oligomerization assay

Performing the cross-linking reaction, the cell lysate was equally separated in two tubes and treated with equal volume of PBS (as a control) or 0.02% glutaraldyhyde for 20 minutes at 25°C. The reaction was stopped by adding 1M Tris buffer (pH8.0) to a final concentration of 50mM Tris. Added 2x laemmli buffer and boiled at 100°C for 15 minutes, followed by western blotting.

### Measuring cell viability

Cells (1.5×10^3^/well) were seeded in 96 well plate for overnight and added drugs for 3 days. Removed medium and added 100 μl (10 μl CCK-8 + 90 μl medium) diluted CCK-8 in each well. Added CCK-8-contained medium into extra well without cells as background control. Incubated at 37°C for 2 hours. Cell viability was measured at OD450/655 nm.

### Colony formation assay

NeHep-SNAP-LHBS cells were seeded in 6 cm dish and drug treatment of adding DMSO (control) and 20 μM TEPP-46 or 25 μM and 50 μM DASA-58 for 2 days. Each 6 cm dish, prepared the bottom agar layer with 1 ml mixture containing ddH_2_O, 2x complete medium, and 2.4% gel (1:2:1). Kept at room temperature becoming solid. Next, trypsinized cells into single cell suspension and counted cell numbers to adjust to 5×10^4^ cells/ml with normal medium. Prepared the top agar layer with 2 ml mixture containing ddH_2_O, 2x complete medium, 2.4% gel (2:1:1), and 2×10^3^ cells in 1 ml normal medium. After top agar gel became solid, added 1.5ml normal medium to provide nutrients. Incubated at 37 °C for 3 weeks and added 1.5 ml complete medium on top of the gel once per week. Dyed the colonies using 10% trypan blue and washed with ddH_2_O, followed by counting colony numbers.

### Extracellular HBsAg and HBeAg Measurement

The supernatant was collected after cell treatments. Centrifuged at full speed for 5 minutes to remove cell debris. HBsAg and HBeAg were measured using Elecsys HBsAg II (cobas) and Elecsys HBeAG (cobas) by cobas e 411 analyzer (Roche).

### 6-Phosphogluconic Acid (6-PGA) Assay

The assay followed the protocol of 6-Phosphogluconic Acid (6-PGA) Assay Kit (Colorimetric) (BioVision). The cells were lysed and centrifuged to get clear extract. Added 50 μl diluted sample into 96 well plate then 50 μl reaction mix for each well. Incubated at 37°C for 1 hour and measured OD 450 nm. The concentration of 6-PGA was calculated against the standard curve.

### Quantification and statistical analysis

All experiments were repeated three times. Western blot was detected by ImageQuant LAS 4000 digital imaging system (GE Healthcare) and quantified using ImageQuant TL software (GE Healthcare). Statistical analysis was performed using Student *t* test with Excel. All data were displayed as mean ± SEM.

## Acknowledgments

We thank Dr. Daniel KY Wu for his personal support.

## Supporting information captions

**S1 Fig. Determine the time of cross-linking using glutaraldehyde (GA).** The cell lysates of control and LHBS-expressing cells were cross-linked for 0, 10, 15, 20, 25, 30 minutes and PKM2 oligomerization were performed by western blot, followed by blotting for PKM2. GA, glutaraldehyde.

**S2 Fig. Infection of Ad-HBV-WT decreases PKM2 kinase activity.** HuH-7 cells were infected with Ad-HBV-KO or Ad-HBV-WT for 2 days and lysates were collected. (A) Expressions of viral LHBS and Tubulin were shown. (B) Pyruvate kinase activity of PKM2 was measured in HuH-7 infected cells. ***, *P*<*0.001*. KO, knockout; WT, wildtype.

